# A pilot study for deciphering post-translational modifications and proteoforms of tau protein by capillary electrophoresis-mass spectrometry

**DOI:** 10.1101/2024.07.04.602093

**Authors:** Fei Fang, Tian Xu, Hsiao-Tien Chien Hagar, Stacy Hovde, Min-Hao Kuo, Liangliang Sun

## Abstract

Abnormal accumulation of tau proteins is one pathological hallmark of Alzheimer□s disease (AD). Many tau protein post-translational modifications (PTMs) are associated with the development of AD, such as phosphorylation, acetylation, and methylation. Therefore, a complete picture of PTM landscape of tau is critical for understanding the molecular mechanisms of AD progression. Here, we offered a pilot study of combining two complementary analytical techniques, capillary zone electrophoresis (CZE)-tandem mass spectrometry (MS/MS) and reversed-phase liquid chromatography (RPLC)-MS/MS, for bottom-up proteomics of recombinant human tau-0N3R. We identified 53 phosphorylation sites of tau-0N3R in total, which is about 30% higher than that from RPLC-MS/MS alone. CZE-MS/MS provided more PTM sites (i.e., phosphorylation) and modified peptides of tau-0N3R than RPLC-MS/MS, and its predicted electrophoretic mobility helped improve the confidence of the identified modified peptides. We developed a highly efficient capillary isoelectric focusing (cIEF)-MS technique to offer a bird’s-eye view of tau-0N3R proteoforms, with 11 putative tau-0N3R proteoforms carrying up to nine phosphorylation sites and lower pI values from more phosphorylated proteoforms detected. Interestingly, under a native-like cIEF-MS condition, we observed three putative tau-0N3R dimers carrying phosphate groups. The findings demonstrate that CE-MS is a valuable analytical technique for the characterization of tau PTMs, proteoforms, and even oligomerization.

## Introduction

Over 20 adult neurodegenerative diseases, such as Alzheimer’s disease (AD), are characterized by the dysfunction and accumulation of tau protein. (1–4) However, the molecular mechanisms that induce tau misfolding and aggregation in neurodegenerative disorders remain elusive. (5) Tau protein post-translational modifications (PTMs) are associated with the development of AD, such as phosphorylation, acetylation, and methylation. (6–9) Therefore, a complete picture of the PTM landscape of tau is critical for understanding the roles played by tau protein in modulating AD progression.

The analytical methods for tau PTM characterization include neuroimaging, biosensing, immunoassay, and mass spectrometry (MS) techniques. (10,11) Among them, immunoassay, which is simple to implement and provides semiquantitative information, has been the workhorse for the identification and quantification of tau or phosphorylated tau (p-tau). (12) However, the antibody-based targeted approach impedes the discovery of new PTMs that could be critical for AD progression.

Emerging MS-based proteomics approaches are powerful alternative methods used to characterize PTMs. In particular, reversed-phase liquid chromatography-tandem MS (RPLC-MS/MS) has been widely used for the characterization of tau PTMs. (13) For example, Steen et al. developed a MS-based assay, Full-Length Expressed stable Isotope-labeled Tau (FLEXITau), with 55 phosphosites, 17 ubiquitination sites, 19 acetylation sites, and 4 methylation sites mapped along 2N4R tau protein from 91 brain tissues. (7,14,15) With targeted parallel reaction monitoring strategy, Barthélemy et al. detected 29 distinct phosphorylated tau sites in the brain and cerebrospinal fluid in humans. (16)

Due to its high separation performance and high detection sensitivity, capillary zone electrophoresis (CZE)-MS/MS is becoming an attractive tool to monitor highly modified samples such as histone proteoforms (17–19) and post-translationally modified peptides. (20) With CZE separating analytes based on their electrophoretic mobilities corresponding to their charge-to-size ratios, not their hydrophobic nature, the combination of RPLC-MS/MS and CZE-MS/MS could impressively expand the PTM coverage. (20,21) However, to the best of our knowledge, no investigators have reported on tau PTM analysis with the combination of CZE-MS/MS and RPLC-MS/MS.

In this work, for the first time, capillary zone electrophoresis (CZE)-MS/MS was investigated for the characterization of tau PTMs and was compared with RPLC-MS/MS in terms of identified modified peptides. We also applied capillary isoelectric focusing (cIEF)-MS to measure the intact tau proteins for a bird’s eye view of tau proteoforms carrying combinations of PTMs.

## Experimental Section

### Chemicals

All chemicals were purchased from Sigma-Aldrich (St. Louis, MO) unless otherwise specified. LC-MS grade solvents, including water, acetonitrile (ACN), methanol, formic acid (FA) and acetic acid were purchased from Fisher Scientific (Pittsburgh, PA). Acrylamide was purchased from Acros Organics (NJ, USA). Fused silica capillaries (50 μm i.d./360 μm o.d.) were purchased from Polymicro Technologies (Phoenix, AZ).

### p-tau (0N3R) expression in E. coli cells and purification

The designing principle of plasmids and procedures for the preparation of PIMAX phosphorylated tau (p-tau) is based on the reference. (8, 22) Briefly, cyclin-dependent kinase 5 (CDK5) and the human 0N3R tau isoform were brought to proximity by heterodimerization of Fos and Jun leucine zipper domains. The overnight culture of BL21-CodonPlus cells harboring PIMAX tau plasmid was diluted to 0.03 OD600 and grown at 37 °C shaking at 230 rpm until reaching 0.3-0.5 OD600. Then, protein expression was induced at 37°C for 2 hours with 0.2 mM IPTG, and the bacterial pellet was collected by centrifugation. The pellet from 1 L induction was suspended with 10 mL lysis buffer (20 mM Tris, pH 5.8, 100 mM NaCl) with 1 mg/mL lysozyme, 1 mM PMSF, 0.2 mM orthovanadate, and 1 tablet of Roche protease inhibitor and incubated for 45 min at room temperature. The suspension was sonicated afterward (Branson Digital Sonifier 450; 30% amplitude; total process time 3 min; pulse-ON time 5 s; pulse-OFF time 5 s). The sonication supernatant was boiled for 30 min in boiling water after the centrifugation at 17,000g, 30 min at 4 °C and the boiled sample was centrifuged afterward. The boiled supernatant was digested with purified recombinant TEV protease in the ratio of 1 OD280 per 100 OD280 of the sample in the presence of 0.5 mM DTT and 1 mM EDTA. The reaction was incubated at 4 °C overnight. The digestion mixture was centrifuged at 17,000g for 30 min at 4 °C, and the supernatant was transferred to another tube and concentrated with Amicon spin column (Amicon Centrifugal Filter Unit, Ultra-15, 30 K) at 5000g at 4 °C. The buffer was changed to storage buffer (20 mM Tris-HCl pH 7.4, 100 mM NaCl) in the meantime. The concentrated sample was supplemented with 10% glycerol (v/v) and stored at -80 °C for further usage.

### Preparation of human p-tau (0N3R) protein for bottom-up proteomics

The protein digestion was performed with the filter-aided sample preparation (FASP) protocol with minor modifications. (23) 20 µg of the p-tau protein sample were loaded onto Microcon 30 kDa molecular weight cutoff centrifugal filter and the buffers were exchanged to 50 µL of 8 M urea dissolved with 50 mM ammonium bicarbonate (NH_4_HCO_3_). The filter unit including the samples was subjected to denaturation and reduction by incubating at 37 °C for 30 minutes with the addition of 10 mM dithiothreitol (DTT). Followed by alkylation with 25 mM iodoacetamide (IAA) for 30 min in the dark, the filter unit was washed with 50 mM NH_4_HCO_3_ four times at 14,000 g. The samples were suspended in 50 µL of 50 mM NH_4_HCO_3_ after clean-up. Afterward, trypsin was added to the sample at a mass ratio of 1:30 for sample digestion. The filter units were further incubated at 37 °C for 4 hours. The peptides from protein digestion were collected by centrifugation at 14,000g for 15 minutes. To reduce the sample loss due to incomplete digestion, the filter units added an additional 20 µL of 50 mM NH_4_HCO_3_ buffer and further incubated at 37 °C overnight. The flow-through was collected by centrifugation and combined with corresponding peptide samples from 4 hours of digestion.

### CZE-MS/MS and RPLC-MS/MS for bottom-up proteomics

The sample digest of p-tau was analyzed by CZE-MS/MS and RPLC-MS/MS, respectively. A CESI 8000 Plus CE system (Sciex) was coupled to Q-Exactive HF mass spectrometer (Thermo Fisher Scientific) via a commercialized electrokinetically pumped sheath-flow CE-MS nanospray interface (EMASS-II, CMP Scientific) for CZE-MS/MS of p-tau peptides. (24,25) A one-meter linear polyacrylamide (LPA)-coated capillary (50 µm I.D., 360 µm O.D.) (26,27) was used for separation and 100 nL of sample was loaded into the capillary at 5 psi for 19 seconds. The glass emitter (orifice size: 30-35 µm) on the interface was filled with sheath liquid (0.2% FA and 10% methanol, pH 2.2) to produce electrospray at 2.2∼2.5 kV. With 5% (v/v) acetic acid used as background electrolyte, 30 kV voltage was applied to the injection end for 110 min, followed by 30 kV voltage and 10 psi pressure applied for 10 min to flush the capillary.

RPLC separation was performed on EASY nanoLC-1200 system (Thermo Fisher Scientific). Mobile phase A comprises 0.1% FA and 5% ACN in water and mobile phase B is 0.1% FA in water/ACN (20:80). With the p-tau digest diluted 10 times with mobile phase A, 2 µL of sample was loaded onto an RPLC capillary column (75 μm I.D., 50 cm length, C18 beads, 2 μm, 100 Å pore size, Thermo Fisher Scientific). RPLC gradient was processed from 2% B to 8% B in 5 minutes, from 8% B to 35% B in 75 minutes, from 35% B to 80% B in 5 minutes, and then stayed at 80% B for 15 minutes at a flow rate of 180 nL/min.

A Q-Exactive HF (Thermo Fisher Scientific) mass spectrometer was used for both CZE-MS/MS and RPLC-MS/MS analyses. The data was collected in DDA mode. For a full MS scan, the resolution was set to 60,000 (at m/z of 200), the m/z range was 300-1500, the AGC target was 3e6, the maximum injection time was 50 ms, and the loop count was 10 (top 10). Fragmentation was performed with an isolation window of 2 m/z. Different normalized HCD energies including 25, 28, and 30 were investigated. MS/MS data was acquired at a resolution of 60,000 (at m/z 200), m/z range of 200-2000, minimal AGC target of 1e4, and maximum injection time of 200 ms. The ion intensity threshold was set to 5e4, and the dynamic exclusion window was 30 s.

### Capillary isoelectric focusing (cIEF)-MS/MS-based top-down analyses of intact p-tau proteoforms

50 μg of purified p-tau protein (0N3R, in 0.1 mM NaCl and 20 mM Tris, pH 7.5) were loaded onto Amicon 10 kDa molecular weight cutoff centrifugal filters and were centrifuged at 14,000g for 15 minutes at 10 °C. The samples were washed four times with 10 mM NH_4_Ac. Eventually, 50 μL of the sample (∼1 mg/mL) was recovered from each filter. For cIEF separation, the samples were further incorporated with ampholytes and glycerol to form a mixture containing 0.5 mg/mL protein, 1.5% ampholyte (narrow-range 8-10.5 and wide-range 3-10 at the ratio of 4:1), and 15% glycerol.

The capillary isoelectric focusing (cIEF) separation was performed on the same CE system for bottom-up analysis using a one-meter LPA-coated capillary. The sheath liquid was 0.2% FA and 10% methanol unless otherwise stated. The capillary was filled with a 6 cm catholyte (0.5% NH_3_·H_2_O, 15% glycerol) at 10 psi for 11 seconds, followed by a 30 cm sample plug at 10 psi for 56 seconds. After sample injection, the capillary inlet was moved to anolyte (0.1% FA, 15% glycerol) with a voltage of 20 kV applied for protein focusing and mobilization. At 20 minutes, an additional pressure of 0.2 psi was applied to assist protein mobilization for 120 min.

One Orbitrap Exploris 480 mass spectrometer (Thermo Fisher Scientific) was used for the cIEF-MS analysis with in-house constructed CE-MS interface. The ion transfer tube temperature of 320 °C, and the RF lens of 60%, the intact protein mode turned on at low-pressure mode. For data-dependent acquisition (DDA), full MS scan was acquired at low resolution of 7,500 (at m/z of 200), m/z range of 600-2500, normalized AGC target of 300%, and microscans of 10. The top 6 most intense precursors (charge states of 5-60, minimal intensity of 1e4) in full MS spectra were selected for fragmentation (isolation window of 2 m/z, HCD energy of 30%). MS/MS data was collected at a resolution of 60,000 (at m/z 200), m/z range of 200-2000, microscans of 10, normalized AGC target of 100%, and auto maximum injection time. The dynamic exclusion was applied with a duration of 30 s and with exclusion of isotopes enabled.

### Data analysis for bottom-up and top-down proteomics

For the bottom-up experiment, the raw files for CZE-MS/MS and RPLC-MS/MS were respectively analyzed by Proteome Discoverer (v 2.2). The raw files were searched against the sequence of human fetal-tau (Uniprot ID P10636-2) and Escherichia coli (E. coli, 4,593 entries, Aug 24, 2023) sequence database downloaded from UniProt (https://www.uniprot.org/). The precursor and fragment ion mass tolerances were 20 ppm and 0.05 Da, respectively. Trypsin was selected as the protease with up to three missed cleavages. The dynamic modification was oxidation (+15.995) on methionine (M), phosphorylation (+79.966) on serine (S)/threonine (T)/tyrosine (Y), acetylation (+42.011) on lysine (K)/arginine (R) and protein N-terminus, succinylation (+100.016) on lysine (K), methylation (+14.016) on lysine (K)/arginine (R) and carbamidomethylation (+57.021) on cysteine (C) was set as static modifications. The target and decoy approach was used to determine the false discovery rates (FDRs) of the identifications. The FDRs were 1% for both peptide and protein. For the modified peptides, only the PTM sites with localization probability higher than 75 were included.

For the top-down experiment, charge state spectra were deconvoluted by the program UniDec version 6.0.4. (28) The charge range is 1 to 100 while the mass range is 5000 to 1000000, with sample mass every 1 Da. Automatic m/z peak width is enabled. The MS2 spectra were processed using ProSight Lite software (29) after deconvolution with TopPIC (top-down mass spectrometry-based proteoform identification and characterization) software (version 1.6.3). (30)

### Experimental and predicted electrophoretic mobility (μ_ef_)

The prediction was performed according to the published literature. (31) The electroosmotic flow (EOF) was assumed zero in the LPA-coated capillary with 5% AA (pH 2.4) as the BGE. For calculating experimental μ_ef_, eq 1 was used,

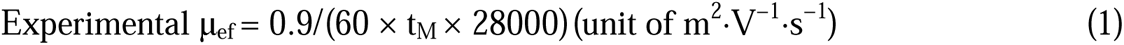

Where migration times (t_M_, min) were determined as time obtained from the database search results. To calculate predicted μ_ef_, eq 2 was used,

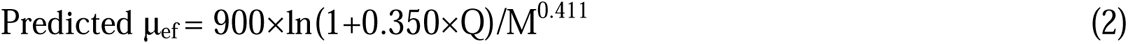

Where Q is the number of charges of each peptide in the liquid phase, represented by the number of positively charged amino acid residues in the peptide sequence (K, R, H, and N-terminus). M is the molecular mass equal to the mass reported by the search engine in Da.

## Results and Discussion

### CZE-MS/MS is an efficient analytical method for bottom-up proteomics of human phospho-tau (p-tau) protein

Due to the high separation efficiency and high detection sensitivity, CZE-MS is a very efficient analytical method for the analysis of peptides and proteins. (19) In this work, we employed both CZE-MS/MS and RPLC-MS/MS for the analysis of tryptic peptides of the shortest recombinant human tau isoform, 0N3R, phosphorylated by cyclin-dependent kinase 5 (CDK5). (32,33) We used *p-tau-0N3R* to represent the CDK5-treated tau 0N3R protein throughout the manuscript. Apart from the most common tau PTM, phosphorylation, a number of other ways, such as glycosylation, acetylation, glycation, ubiquitylation, *O*-GlcNAcylation, aggregation, and filament formation can be modified on tau protein in neurodegenerative disease brain. (34) While the current p-tau was synthesized in E. coli for the creation of human AD-relevant hyperphosphorylation pattern (8, 22), additional PTMs such as methylation, acetylation (35) and succinylation (36,37) that are known to occur in many recombinant proteins produced in *E. coli* were also present in this and other PIMAX p-tau preparations (see below, and unpublished observations by Hagar and Kuo).

As a result, RPLC-MS/MS identified peptides covering 94% of the *p-tau-0N3R* sequence (**Figure S1A**) while CZE-MS/MS achieved 100% sequence coverage (**Figure S1B**). Among them, only 66% of the peptides carrying PTM sites were commonly identified with RPLC-MS/MS and CZE-MS/MS, showing the different peptide identification preferences for these two methods (**Figure 1A**). As shown in **Figure 1B**, both methods can identify peptides with multiple PTM sites, with up to four occurrences per peptide. In addition, the CZE-MS/MS could identify more peptides carrying phosphate group or succinyl group (**Figure 1C**). The different peptide sequences identified from RPLC-MS/MS and CZE-MS/MS demonstrate the high complementarity of these two methods for tau PTM analysis. The lists of peptides identified from tau-0N3R are listed in **Supporting Information II**.

**Figure 1.**
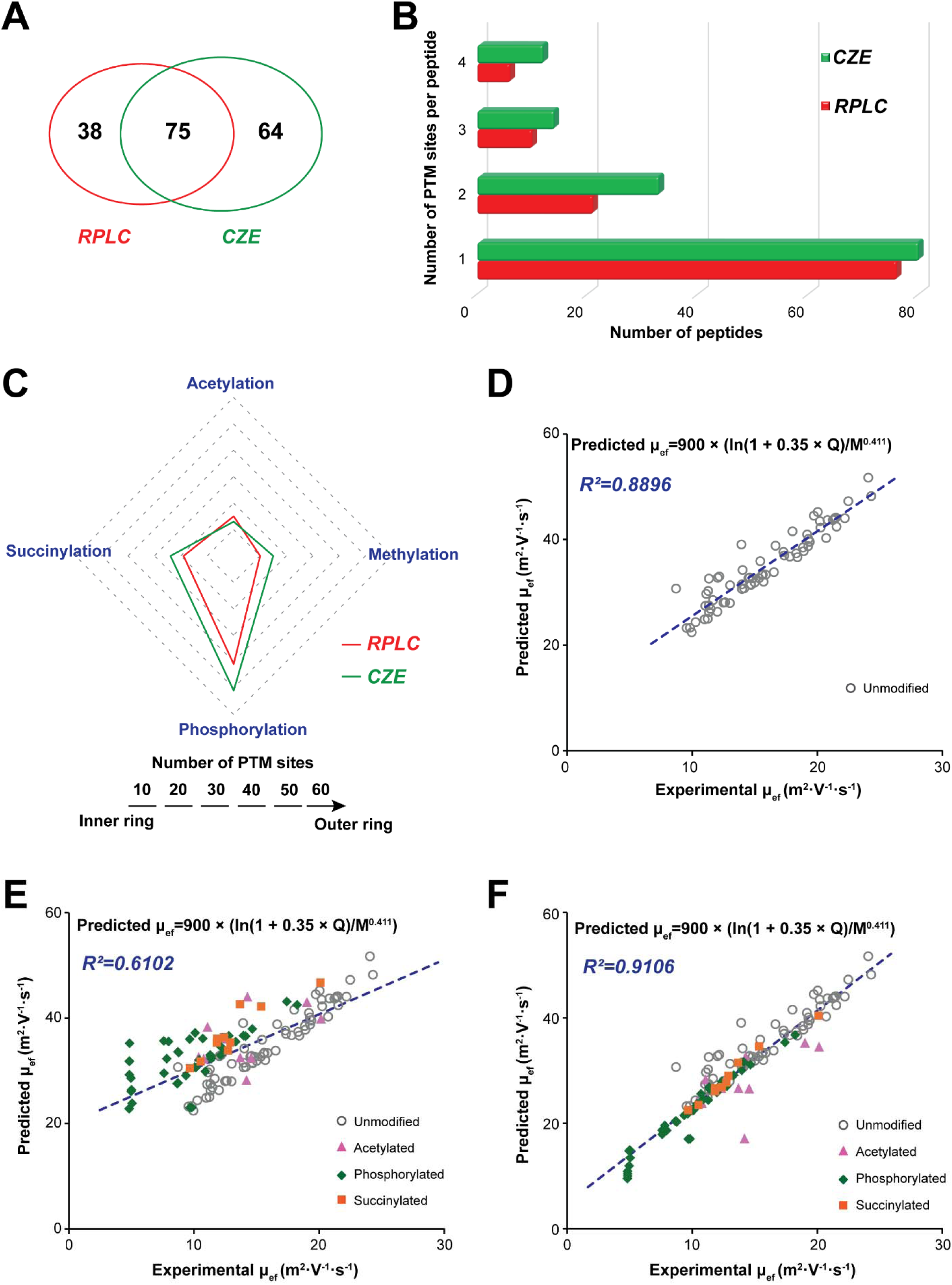
CZE-MS/MS and RPLC-MS/MS analysis of tryptic digest from human full-length human phosphorylated tau-0N3R protein (*p-tau-0N3R*) protein. The comparison of (A) number of peptides with PTM, (B) number of PTM sites per peptide and (C) number of PTM sites identified with RPLC-MS/MS and CZE-MS/MS. Linear correlations between theoretical and experimental µ_ef_ of (D) unmodified peptides and (E) unmodified plus phosphorylated, acetylated and succinylated peptides identified from p-tau sample with CZE-MS/MS analysis under the BGE of 5% (v/v) acetic acid (pH 2.4). (F) Linear correlations between theoretical and experimental µ_ef_ of unmodified peptides plus phosphorylated, acetylated and succinylated peptides after charge Q corrections due to PTMs. In Figure F, we adjusted the Q by -1, -1, and -1 for peptides with phosphorylation, acetylation, and succinylation, respectively. The grey circles represent peptides without PTMs, the purple triangles represent acetylated peptides, the green diamonds represent phosphorylated peptides, and orange squares represent succinylated peptides.

We applied a multiparametric sequence-specific model (38) to predict the electrophoretic mobility for the peptides identified with CZE-MS/MS, which could help validate the confidence of peptide identifications by examining the correlation between predicted and experimental µ_ef_ of peptides. In the Experimental Section, we described the details of calculating the predicted and experimental µ_ef_ of the peptides.

Firstly, the peptides without any PTMs produced a linear correlation coefficient (R^2^) of 0.8896, **Figure 1D**. Then, the µ_ef_ prediction equation was used for p-tau peptides with PTMs (acetylation, phosphorylation, and succinylation). Since those PTMs usually lead to reduced mobility of peptides, most of the p-tau peptides with PTMs were clearly off the trendline without charge Q corrections, **Figure 1E**. Peptide acetylation and phosphorylation were reported to reduce the charge Q by roughly one unit according to our previous studies. (38) According to the chemical reaction involved in succinylation, we expect that one succinylation reduces the peptide charge Q by one unit in our pH 2.4 BGE condition. After we applied charge Q reduction for those acetylated, phosphorylated, and succinylated peptides, the linear correlation coefficient between predicted and experimental µ_ef_ increased from 0.6102 (**Figure 1E**) to 0.9106 (**Figure 1F**). The data demonstrate that µ_ef_ of modified tau peptides can be predicted accurately and that one succinylation PTM certainly reduced the peptide charge Q by one unit in our CZE-MS condition. An example of the peptide carrying 1, 2 or 3 phosphate groups identified with CZE-MS/MS is shown in **Figure S2**. Peptides that carry more phosphate groups migrate more slowly. As illustrated in **Figure S3**, in comparison with the unmodified peptide, the peptides with the same sequence but carrying one phosphate group or succinyl group were eluted later at almost the same time, which agrees well with the prediction result.

We combined all the PTM sites identified with RPLC-MS/MS and CZE-MS/MS (**Figure 2**). Interestingly, it was found that more PTM sites located in the proline-rich domain were discovered by CZE-MS/MS, while RPLC-MS/MS favored the identification of PTM sites of the C-terminal half, further demonstrating the high complementary between these two methods for PTM identification of tau. The PTM mapping showed that succinylation occurred widely along the tau sequence, and lysines that could be methylated or acetylated are also frequent targets of succinylation, which agrees well with the literature. (39)

**Figure 2.**
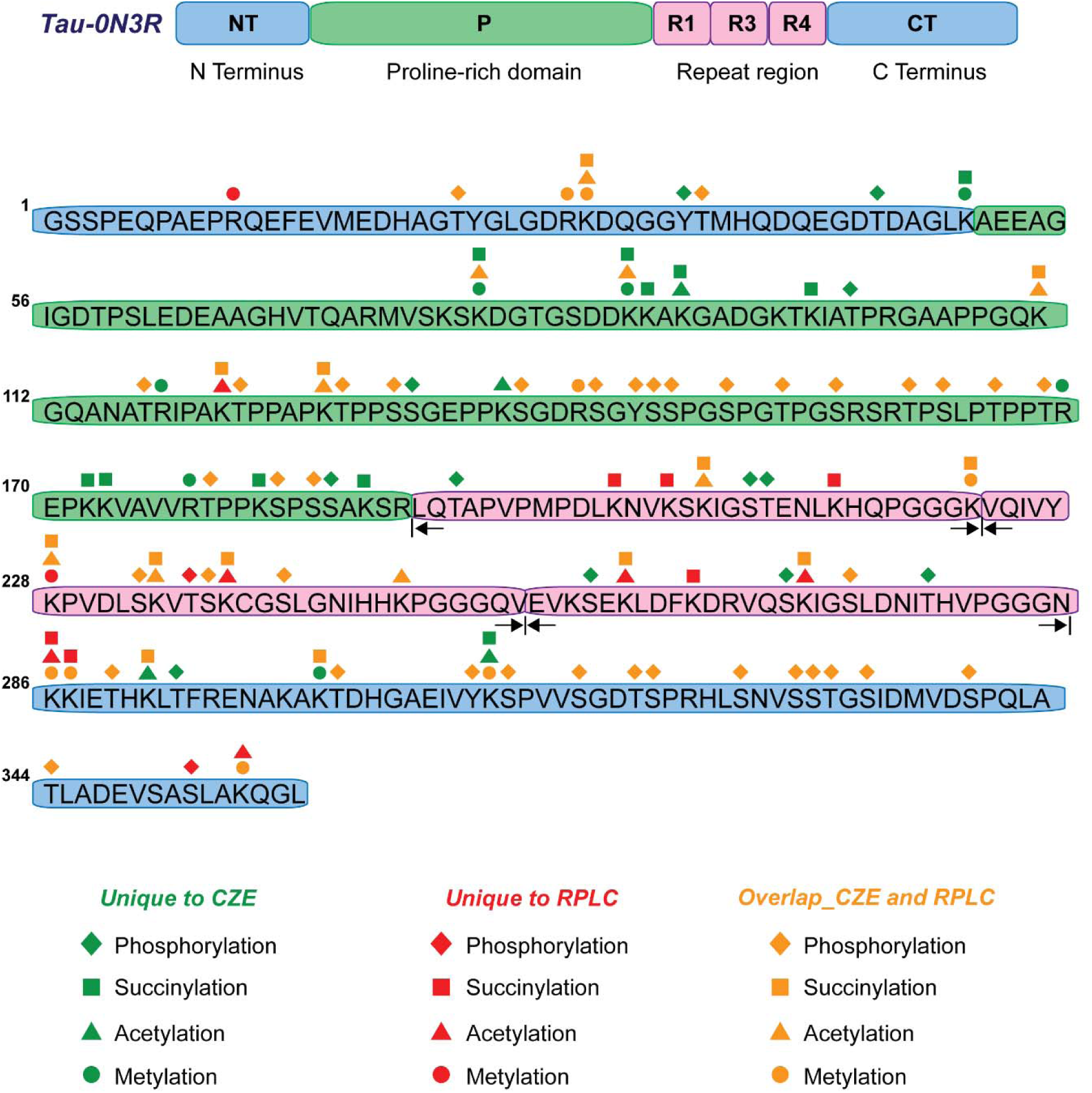
Bar diagram of human *p-tau-0N3R* with the distribution of observed PTM sites. The diamond, square, triangle, and circle represent the possible phosphorylation, succinylation, acetylation, and methylation sites, respectively, identified from bottom-up proteomics. The green and red symbols represent the PTM sites uniquely identified by CZE-MS/MS and RPLC-MS/MS, respectively. The orange symbols represent the PTM sites commonly identified by CZE-MS/MS and RPLC-MS/MS.

Among the 67 putative phosphorylation sites on *p-tau-0N3R*, the number of P-sites identified on serine (S), threonine (T) and tyrosine (Y) residues on tau was 41 sites (23 S, 16 T, and 2 Y) from RPLC-MS/MS and 51 sites (27 S, 21 T and 3 Y) with CZE-MS/MS (**Table S1**). The total number of 53 detected phosphorylated residues represents 53/67 = 79% of all potential phosphorylated residues, which occupied almost all the potential phosphorylation sites in the repeat region (11/13) and C-terminal domains (16/17).

Hanger et al. compiled the phosphorylation sites for tau protein, with 12 phosphorylation sites detected from the CDK5 phosphorylated tau protein. (40) Kuo and colleagues applied four endoproteinases, trypsin, Lys-C, Arg-C or Asp-N for in-gel digestion followed by RPLC-MS/M and identified 31 phosphorylation sites from human tau-0N3R. (9) As shown in **Table S2**, all phosphorylation sites observed by Hanger et al. were detected in our work. In comparison with Kuo’s result, most of the phosphorylation sites, except Thr-53 and Thr-91, were identified by our method. Among the 24 phosphorylation sites uniquely identified in our result, 8 of them were uniquely identified with CZE-MS/MS method, which further indicated the high promise of CZE-MS/MS for a global characterization of tau PTMs.

### cIEF-MS analysis of intact human p-tau-0N3R proteoforms

Our bottom-up proteomics measurement reveals rich PTM information (e.g., 53 phosphorylation sites) of human *p-tau-0N3R*. However, we cannot figure out the pictures of intact *p-tau-0N3R* proteoforms based on the bottom-up proteomics data. For example, what types of *p-tau-0N3R* proteoforms are there? How different PTMs are organized on each *p-tau-0N3R* proteoform? Top-down proteomics measurement of tau proteins is critical for filling this knowledge gap and is fundamental for understanding the roles of tau protein in AD progression because tau functions as intact proteoforms carrying various combinations of PTMs, not individual peptides carrying PTMs. Therefore, in this work, we also investigated CE-MS for top-down proteomics analysis of human *p-tau-0N3R*.

Top-down proteomics has been tested for characterizing intact tau proteoforms in the literature. (10,16) Mandelkow *et al.* analyzed the intact phosphorylated tau (2N4R) expressed in eukaryotic cells by direct infusion native MS, revealing the distribution of the number of phosphates per tau protein. (10) Loo *et al.* integrated ion mobility spectrometry and electron capture dissociation with direct infusion native MS for the intact tau and tau/CLR01 complex to pinpoint the site(s) of inhibitor binding. (16) Online liquid-phase high-resolution tau proteoform separation prior to MS is essential for the detection of low-abundance tau proteoforms and the generation of a complete tau proteoform landscape.

In this work, we first tested CZE-MS for the characterization of *p-tau-0N3R* proteoforms. However, we did not observe clear intact tau proteoform signals, maybe due to the high complexity the *p-Tau-0N3R* proteoforms and coelution with other background proteins. We decided to investigate cIEF-MS for this purpose due to the super high separation resolution of cIEF for proteins/proteoforms with isoelectric point (pI) difference and its much higher sample loading capacity, which could be critical for highly complex and large proteoforms. We performed cIEF-MS analysis of *p-tau-0N3R* with about 300 ng tau protein injected and detected 11 potential p-tau-0N3R proteoforms after mass deconvolution with UniDec software within 9 electrophoretic peaks, **Figure 3A**. The mass of detected potential 0N3R proteoforms is in a range of 37.6 and 38.2 kDa. The average mass of tau-0N3R proteoform with N-terminal methionine removal and a seven amino acid residue remnant at the N-terminus from cloning (GSSPEQP) and without any PTMs is 37,312 Da. Therefore, the proteoform in P1 (37,771 Da), might represent a tau-0N3R proteoform containing 5 phosphate groups, 1 acetyl and 1 methyl group. The proteoforms in P2 (37,852 Da) and P3 (37,932 Da), show an around 80-Da mass difference with P1, might carrying 6 and 7 phosphate groups, respectively. The original mass spectra and deconvoluted masses are shown in **Figure S4A-C**.

**Figure 3.**
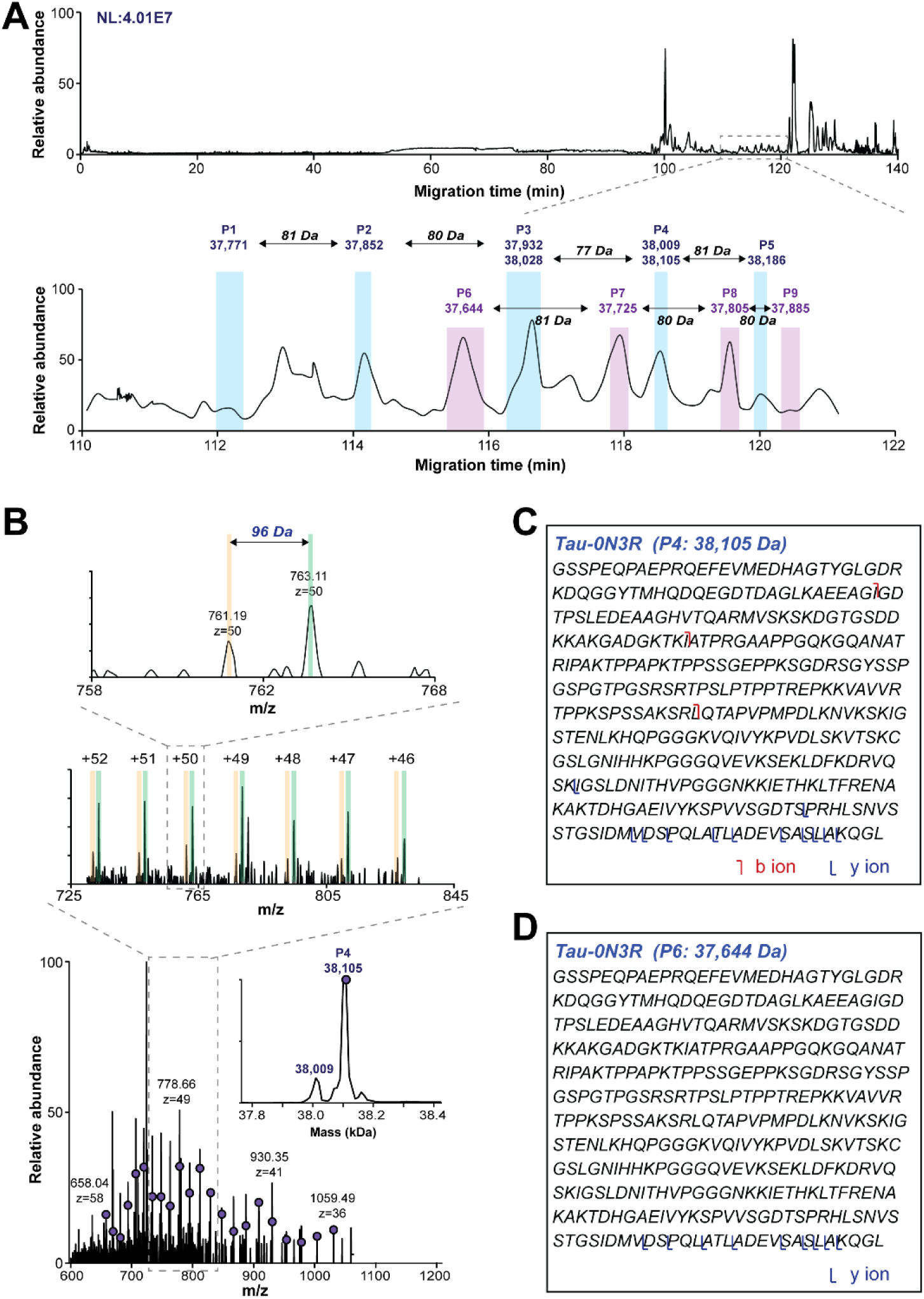
cIEF-MS analysis of human *p-tau-0N3R* under a denaturing condition. (A) Base peak electropherogram of *p-tau-0N3R* proteoforms (top). Zoomed-in base peak electropherogram in 110-122 min with deconvoluted masses of *p-tau-0N3R* proteoforms labeled. (B) Averaged mass spectra and deconvoluted mass spectra (inserted figure) of *p-tau-0N3R* proteoforms detected in peak 4 (P4). A zoomed-in mass spectrum of *p-tau-0N3R* proteoforms was detected in P4. The sequence and fragmentation pattern of *p-tau-0N3R* proteoforms detected in P4 (C) and P6 (D).

We detected another proteoform with 38,028 Da in P3, with a +96 Da mass shift in comparison with the proteoform of 37,932 Da, **Figure S4C**. We also observed two proteoforms in peak 4 with masses of 38,009 and 38,105 Da, **Figure 3B**. Interestingly, the mass difference between 37,932 Da in peak 3 and 38,009 Da in peak 4 is close to 80 Da, and a similar mass difference was also observed for the other two proteoforms in peaks 3 and 4 (i.e., 38,028 vs. 38,105 Da). Peak 5 contains one proteoform (38,186 Da, **Figure S4D**) close to the mass of 0N3R, which is 81 Da heavier than one proteoform in peak 4 (38,105 Da). The data suggests that the tau-0N3R proteoforms in peaks 4 and 5 should carry 8 and 9 phosphate groups. An observation from the data is that the *p-tau-0N3R* proteoform carrying more phosphate groups migrated more slowly due to the reduction of the isoelectric point (pI). Also, the migration time reduction by adding one phosphate group is substantial (i.e., about 2 minutes), **Figure 3A**. Based on the results, we speculate that the 96-Da mass shift between the two proteoforms in peak 3 or peak 4 should not correspond to phosphorylation PTM because they nearly co-migrated. We also detected peak series corresponding to another four *p-tau-0N3R* proteoforms (peaks 6-9) with mass from 37,644 to 37,885 Da with equally spaced by ∼80 Da, **Figure 3A** and **Figure S4 E-H**. The proteoform in P6 (37,644 Da) might be a tau-0N3R proteoform carrying 4 phosphate groups and 1 methyl group, while those proteoforms from P7 to P9 also might be from *p-tau-0N3R* with an increased amount of phosphorylation. We further analyzed the MS/MS data of peaks 4 and 6 and did targeted analysis using ProSight Lite (29), **Figures 3C** and **3D**. A series of fragment ions of those two peaks matched the C-terminus of tau-0N3R, which agrees well with the manual annotation (**Figure S5**). The peaks corresponding to the neutral loss of phosphoric acid shown in **Figure S5** further confirmed those proteoforms are from phosphorylated tau-0N3R. However, we cannot localize the PTM sites on the tau proteoforms due to the high positional heterogeneity of the phosphorylation modifications and the limited fragmentation efficiency of HCD on those large proteoforms. Overall, the results demonstrate that cIEF-MS is a promising technique for the characterization of tau proteoforms with only nanograms of sample consumption. More work needs to be done to improve the backbone cleavage coverage of tau proteoforms for better PTM localization in future studies.

### cIEF-MS analysis of intact human p-tau 0N3R protein under the pseudo-native condition

We also performed a top-down MS analysis of *p-tau-0N3R* under a pseudo-native condition by changing the sheath liquid from 10% methanol and 0.2% FA to 10 mM ammonium acetate (pH adjusted to 4 with acetic acid). We have demonstrated native cIEF-MS for the characterization of protein complexes and antibody-drug conjugates in one of our recent studies. (41) Here, we applied a similar condition to the *p-tau-0N3R*, aiming to achieve more information about the tau protein sample. As expected, with the increase in pH of the sheath liquid, the most abundant charge state of the detected *p-tau-0N3R* proteoforms shifted from around +50 (**Figures S4**) to around +40 (**Figure S6**). We detected some of the same proteoforms as the denaturing condition. For example, the proteoforms in P4, P5, P6, P7, and P8 in **Figure 4A** are the same as those in P1, P2, P6, P3, and P7 in **Figure 3A**.

**Figure 4.**
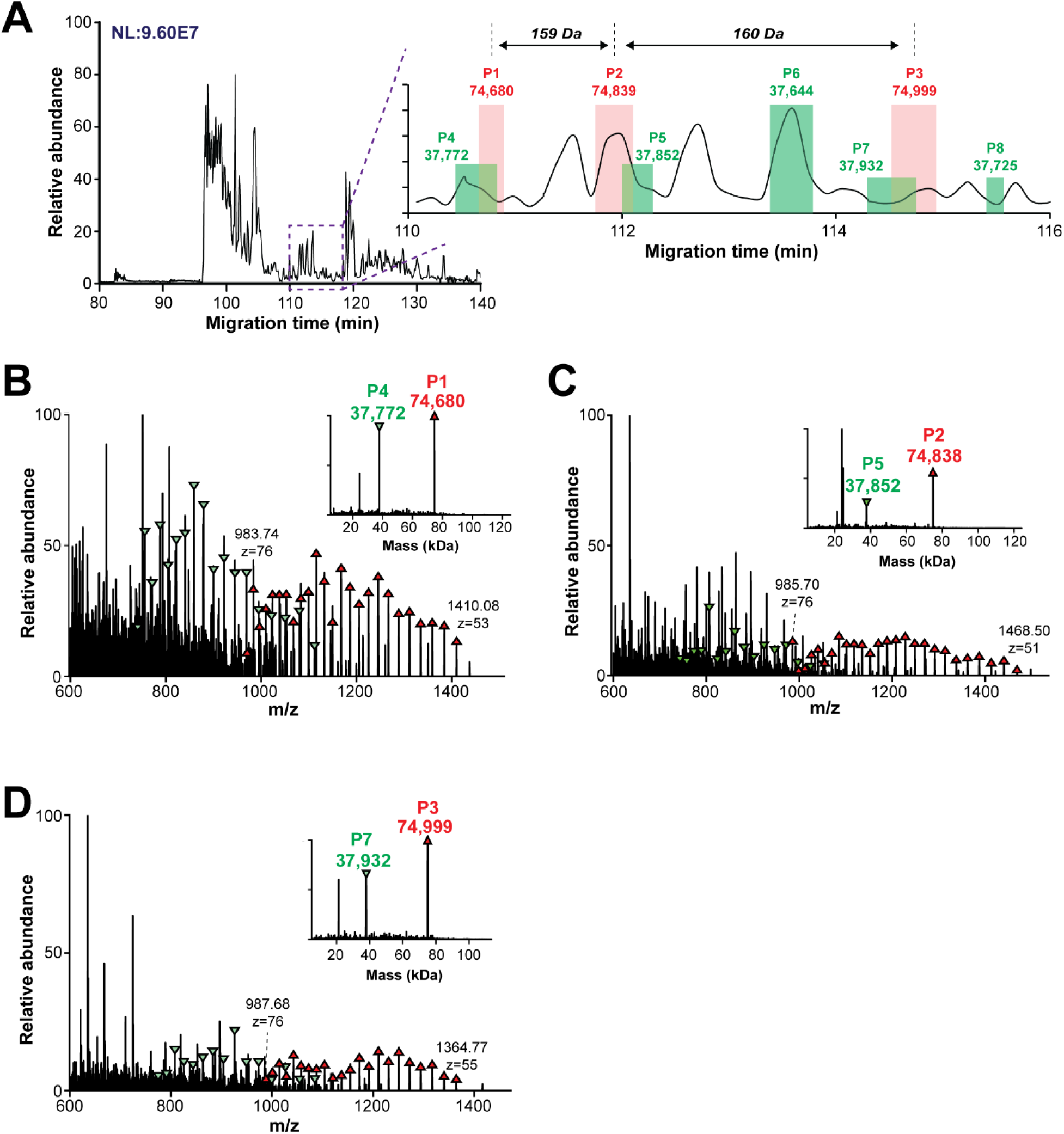
cIEF-MS analysis of human *p-tau-0N3R* under a pseudo-native condition. (A) Base peak electropherogram of *p-tau-0N3R* proteoforms with a zoomed-in base peak electropherogram of the 110-116 min region (inserted figure). The deconvoluted masses of different tau proteoforms are labeled. (B-D) Averaged mass spectra and deconvoluted masses (inserted figures) of *p-tau-0N3R* proteoforms detected in P1-P3.

Interestingly, the pseudo-native condition also resulted in the detection of three large proteoforms ranging from 74,680 to 74,999 Da with equally spaced by ∼160 Da, **Figure 4A**. We speculate that those over 74 kDa proteoforms are dimers of different *p-tau-0N3R* proteoforms. Considering the partial co-migration of proteoforms in P1 and P4, P2 and P5, and P3 and P7, a possible explanation is that the proteoforms in P4, P5, and P7 might contribute to the formation of dimers in P1, P2, and P3. For example, the proteoform dimer of 74,680 Da detected in P1 might be formed by the two tau-0N3R proteoform of 37,772 (tau-0N3R proteoform carrying 5 phosphate groups) with an unknown -864 Da mass shift. Similarly, the large proteoforms detected in P2 and P3 are the dimers carrying 6 and 7 phosphate groups, respectively. These results are in agreement with that the pathological tau protein could form dimers. (42,43) Although the assumption needs to be confirmed with a more efficient fragmentation method for better tau proteoform characterization, it demonstrates the high promise of pseudo-native cIEF-MS/MS for the top-down MS analysis of specific tau proteoforms in their aggregated forms. Importantly, our preliminary data suggests that the dimer formation of *p-tau-0N3R* proteoforms may be selective because proteoforms in P4, P5, and P7 are involved but P6 and P8 are not involved.

## 4. Conclusions

In this pilot study, for the first time, CE-MS/MS was applied for the bottom-up proteomics and top-down proteomics analyses of the human *p-tau-0N3R* expressed in *E. coli* cells. CZE-MS/MS-based bottom-up proteomics analysis mapped 51 phosphorylation sites,13 acetylation sites, 15 methylation sites, and 24 succinylation sites. CZE-MS/MS provided more PTM sites of *p-tau-0N3R* than RPLC-MS/MS and the two techniques showed nice complementarity regarding tau PTM characterization. CIEF-MS/MS-based top-down analysis of *p-tau-0N3R* proteoforms offered a bird’s-eye view of tau proteoforms in the sample with high sensitivity. The pseudo-native cIEF-MS revealed the likely formation of 0N3R dimers from specific proteoforms. The results show that CZE-MS/MS and cIEF-MS/MS are very useful analytical tools for elucidating tau PTMs and proteoforms. It is conceivable that tau extracted from the human brain with varying degrees of Alzheimer’s disease or other tauopathies can be decoded to details by methods described in this work, hence promoting our understanding of the mechanism and possibly the clinical management of these devastating neurodegenerative disorders.

## Acknowledgment

We thank Kuang-Wei Wang for technical assistance, and the financial support from the National Institute of General Medical Sciences (NIGMS) through grant R35GM153479 (Sun) and the National Cancer Institute (NCI) through grant R01CA247863 (Sun). MHK is partly supported by NIH grants R01NS135693, RF1AG077475, and R01AG062435.

## Notes

The authors declare no competing financial interest.

## Data availability

MS raw data and result files have been deposited to the ProteomeXchange Consortium via the PRIDE repository (44) and are publicly accessible from its website with the dataset identifier PXD053367.

## Supporting information

The phosphorylation sites of human *p-tau-0N3R* protein identified by RPLC-MS/MS and CZE-MS/MS with localization probability higher than 75 (Table S1); P-sites in human tau (0N3R) expressed in E. coli cells by three independent studies (Table S2); Bottom-up analysis of phosphorylated human *p-tau-0N3R* protein via RPLC-MS and CZE-MS/MS (Figure S1); Example spectra of peptides carrying different numbers of phosphate groups (Figure S2); Example spectrum of peptide (KDQGGYTMHQDQEGDTDAGLK) without PTMs and with phosphorylation or succinylation (Figure S3); cIEF-MS analysis of human *p-tau-0N3R* under denaturing condition (Figure S4); Spectrum of two human *p-tau-0N3R* proteoforms carrying different numbers of phosphate groups under denaturing conditions (Figure S5); cIEF-MS analysis of human *p-tau-0N3R* under pseudo-native condition (Figure S6) (PDF)

Lists of tryptic peptides identified from human *p-tau-0N3R* by RPLC-MS/MS and CZE-MS/MS (XLSX)

## For TOC only

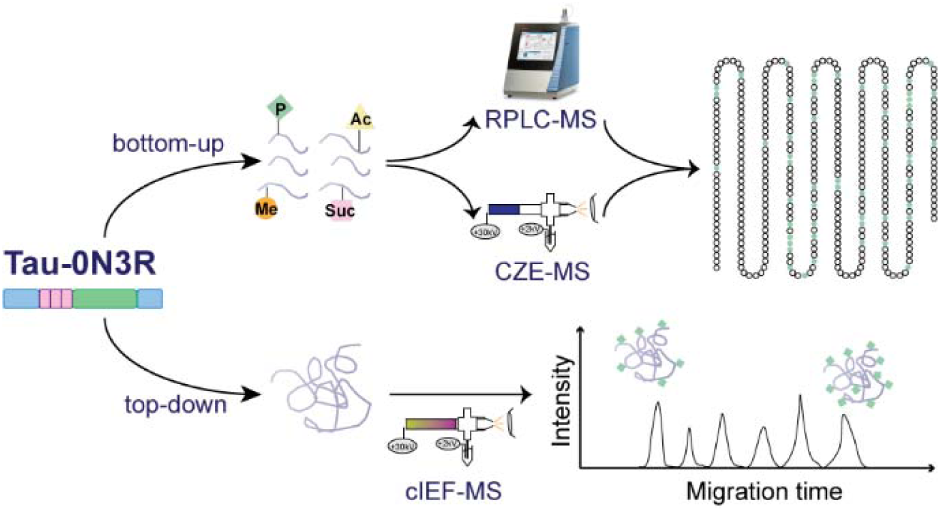

## Notes

### Competing Interest Statement

The authors have declared no competing interest.

