## supporting information I for "A pilot study for deciphering post-translational modifications and proteoforms of tau protein by capillary electrophoresis-mass spectrometry"

**Table of Contents**

| **Supplementary Tables and Figures** | **Page** |
| --- | --- |
| Table S1 | **S3-S4** |
| Table S2 | **S5-S6** |
| Figure S1 | **S7** |
| Figure S2 | **S8** |
| Figure S3 | **S9** |
| Figure S4 | **S10** |
| Figure S5 | **S11** |
| Figure S6 | **S12** |

**Table S1.** The phosphorylation sites of *p-tau-0N3R* protein identified by RPLC-MS/MS and CZE-MS/MS with localization probability higher than 75.

| **Peptide**  **No.** | **Residues** | **P-Site** | **Peptide sequence** | **RPLC** | **CZE** |
| --- | --- | --- | --- | --- | --- |
| 1 | 6-23 | T17 | QEFEVMEDHAG**T**YGLGDR | + | + |
| 2 | 25-44 | Y29 | DQGG**Y**TMHQDQEGDTDAGLK |  | + |
| 3 | 25-44 | T30 | DQGGY**T**MHQDQEGDTDAGLK | + | + |
| 4 | 25-44 | T39 | DQGGYTMHQDQEGD**T**DAGLK |  | + |
| 5 | 91-97 | T95 | TKIA**T**PR |  | + |
| 6 | 106-116 | T111 | GQANA**T**RIPAK | + | + |
| 7 | 113-122 | T117 | IPAK**T**PPAPK | + | + |
| 8 | 123-132 | T123 | **T**PPSSGEPPK | + | + |
| 9 | 123-132 | S126 | TPP**S**SGEPPK | + | + |
| 10 | 123-151 | S127 | TPPS**S**GEPPKSGDRSGYSSPGSPGTPGSR |  | + |
| 11 | 123-151 | S133 | TPPSSGEPPK**S**GDRSGYSSPGSPGTPGSR | + | + |
| 12 | 123-151 | S137 | TPPSSGEPPKSGDR**S**GYSSPGSPGTPGSR | + | + |
| 13 | 133-151 | Y139 | SGDRSG**Y**SSPGSPGTPGSR | + | + |
| 14 | 137-151 | S140 | SGY**S**SPGSPGTPGSR | + | + |
| 15 | 123-151 | S141 | TPPSSGEPPKSGDRSGYS**S**PGSPGTPGSR | + | + |
| 16 | 123-151 | S144 | TPPSSGEPPKSGDRSGYSSPG**S**PGTPGSR | + | + |
| 17 | 137-151 | T147 | SGYSSPGSPG**T**PGSR | + | + |
| 18 | 137-151 | S150 | SGYSSPGSPGTPG**S**R | + | + |
| 19 | 152-163 | T154 | SR**T**PSLPTPPTR | + | + |
| 20 | 154-163 | S156 | TP**S**LPTPPTR | + | + |
| 21 | 154-163 | T159 | TPSLP**T**PPTR | + | + |
| 22 | 154-166 | T162 | TPSLPTPP**T**REPK | + | + |
| 23 | 167-182 | T173 | KVAVVR**T**PPKSPSSAK | + | + |
| 24 | 173-182 | S177 | TPPK**S**PSSA | + | + |
| 25 | 173-182 | S179 | TPPKSP**S**SAK | + | + |
| 26 | 177-182 | S180 | SPS**S**AK |  | + |
| 27 | 185-196 | T187 | LQ**T**APVPMPDLK |  | + |
| 28 | 202-209 | S204 | IG**S**TENLK |  | + |
| 29 | 202-209 | T205 | IGS**T**ENLK |  | + |
| 30 | 217-232 | S227 | VQIVYKPVDL**S**KVTSK | + | + |
| 31 | 217-232 | T230 | VQIVYKPVDLSKV**T**SK | + |  |
| 32 | 217-232 | S231 | VQIVYKPVDLSKVT**S**K | + | + |
| 33 | 217-251 | S235 | VQIVYKPVDLSKVTSKCG**S**LGNIHHKPGGGQVEVK | + | + |
| 34 | 252-260 | S252 | **S**EKLDFKDR |  | + |
| 35 | 261-280 | S263 | VQ**S**KIGSLDNITHVPGGGNK |  | + |
| 36 | 261-280 | S267 | VQSKIG**S**LDNITHVPGGGNK | + | + |
| 37 | 265-280 | T272 | IGSLDNI**T**HVPGGGNK |  | + |
| 38 | 281-290 | T284 | KIE**T**HKLTFR | + | + |
| 39 | 281-294 | T288 | KIETHKL**T**FRENAK |  | + |
| 40 | 295-317 | T297 | AK**T**DHGAEIVYKSPVVSGDTSPR | + | + |
| 41 | 295-317 | Y305 | AKTDHGAEIV**Y**KSPVVSGDTSPR | + | + |
| 42 | 295-317 | S307 | AKTDHGAEIVYK**S**PVVSGDTSPR | + | + |
| 43 | 307-317 | S311 | SPVV**S**GDTSPR | + | + |
| 44 | 307-317 | T314 | SPVVSGD**T**SPR | + | + |
| 45 | 307-317 | S315 | SPVVSGDT**S**PR | + | + |
| 46 | 318-349 | S320 | HL**S**NVSSTGSIDMVDSPQLATLADEVSASLAK | + | + |
| 47 | 318-349 | S323 | HLSNV**S**STGSIDMVDSPQLATLADEVSASLAK | + | + |
| 48 | 318-349 | S324 | HLSNVS**S**TGSIDMVDSPQLATLADEVSASLAK | + | + |
| 49 | 318-349 | T325 | HLSNVSS**T**GSIDMVDSPQLATLADEVSASLAK | + | + |
| 50 | 307-349 | S327 | SPVVSGDTSPRHLSNVSSTG**S**IDMVDSPQLATLADEVSASLAK | + | + |
| 51 | 318-349 | S333 | HLSNVSSTGSIDMVD**S**PQLATLADEVSASLAK | + | + |
| 52 | 318-349 | T338 | HLSNVSSTGSIDMVDSPQLA**T**LADEVSASLAK | + | + |
| 53 | 318-349 | S346 | HLSNVSSTGSIDMVDSPQLATLADEVSA**S**LAK | + |  |

**Table S2.** P-sites in human tau (0N3R) expressed in *E. coli* cells by three independent studies: Hanger et al. (2020) [compile], Kuo et al. (2020) [RPLC-MS/MS] and current work [CZE-MS/MS and RPLC-MS/MS]. Column 1, potential P-sites in full-length human tau (0N3R, residues S, T, Y); column 2, P-sites in human tau compiled by Hanger, 2020; columns 3, P-sites in tau observed in Kuo, 2020; column 4, P-sites in tau observed in current work. Additional sites observed in the current work are marked in red, and sites observed in previous but not in the present study are in blue.

| **Potential P-sites**  **in hTau 0N3R** | **P-sites obs.**  **(Hanger, 2020)** | **P-sites obs.**  **(Kuo 2020)** | **P-sites obs.**  **(this work)** |
| --- | --- | --- | --- |
| T17 |  |  | T17 |
| Y18 |  |  |  |
| Y29 |  |  | Y29 |
| T30 |  |  | T30 |
| T39 |  |  | T39 |
| T53 |  | T53 |  |
| S55 |  |  |  |
| T65 |  |  |  |
| S71 |  |  |  |
| S73 |  |  |  |
| T77 |  |  |  |
| S79 |  |  |  |
| T91 |  | T91 |  |
| T95 | T95 | T95 | T95 |
| T111 |  | T111 | T111 |
| T117 |  | T117 | T117 |
| T123 | T123 | T123 | T123 |
| S126 |  |  | S126 |
| S127 |  |  | S127 |
| S133 |  | S133 | S133 |
| S137 | S137 | S137 | S137 |
| Y139 |  |  | Y139 |
| S140 |  |  | S140 |
| S141 | S141 | S141 | S141 |
| S144 | S144 | S144 | S144 |
| T147 | T147 | T147 | T147 |
| S150 |  |  | S150 |
| S152 |  |  |  |
| T154 | T154 | T154 | T154 |
| S156 | S156 |  | S156 |
| T159 |  | T159 | T159 |
| T162 |  | T162 | T162 |
| T173 | T173 | T173 | T173 |
| S177 | S177 | S177 | S177 |
| S179 |  |  | S179 |
| S180 |  |  | S180 |
| S183 |  |  |  |
| T187 |  | T187 | T187 |
| S200 |  |  |  |
| S204 |  |  | S204 |
| T205 |  |  | S205 |
| Y221 |  |  |  |
| S227 |  | S227 | S227 |
| T230 |  |  | T230 |
| S231 |  | S231 | S231 |
| S235 |  | S235 | S235 |
| S252 |  |  | S252 |
| S263 |  | S263 | S263 |
| S267 |  | S267 | S267 |
| T272 |  | T272 | T272 |
| T284 |  | T284 | T284 |
| T288 |  |  | T288 |
| T297 |  |  | T297 |
| Y305 |  |  | Y305 |
| S307 | S307 | S307 | S307 |
| S311 |  | S311 | S311 |
| T314 |  | T314 | T314 |
| S315 | S315 | S315 | S315 |
| S320 |  | S320 | S320 |
| S323 |  |  | S323 |
| S324 |  |  | S324 |
| T325 |  |  | T325 |
| S327 |  | S327 | S327 |
| S333 |  | S333 | S333 |
| T338 |  |  | T338 |
| S344 |  |  |  |
| S346 |  |  | S346 |

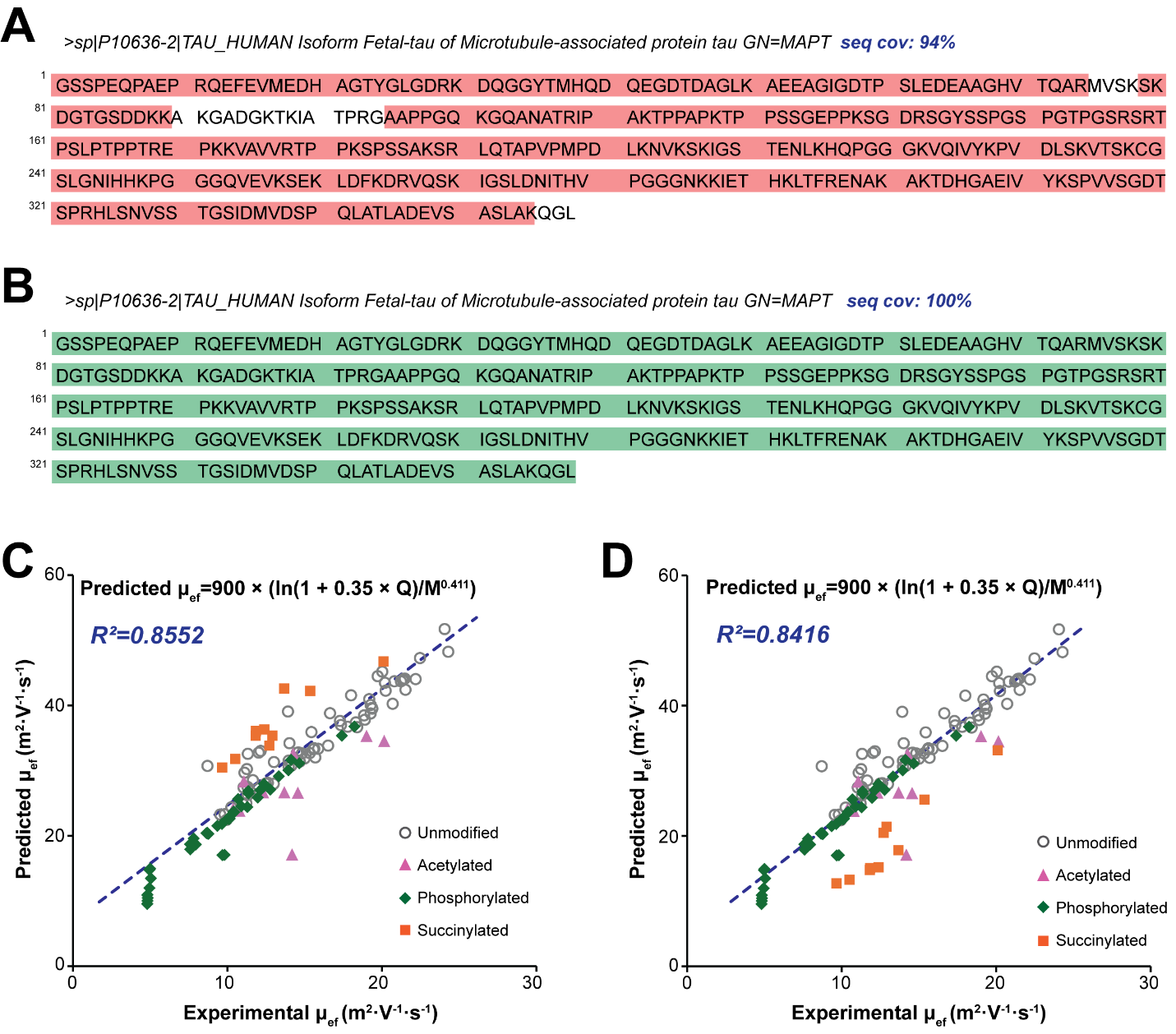

**Figure S1.** Bottom-up analysis of phosphorylated human *p-tau-0N3R* protein via RPLC-MS and CZE-MS/MS. Coverage map of 0N3R isoform of human tau identified by (A) RPLC-MS/MS and (B) CZE-MS/MS.

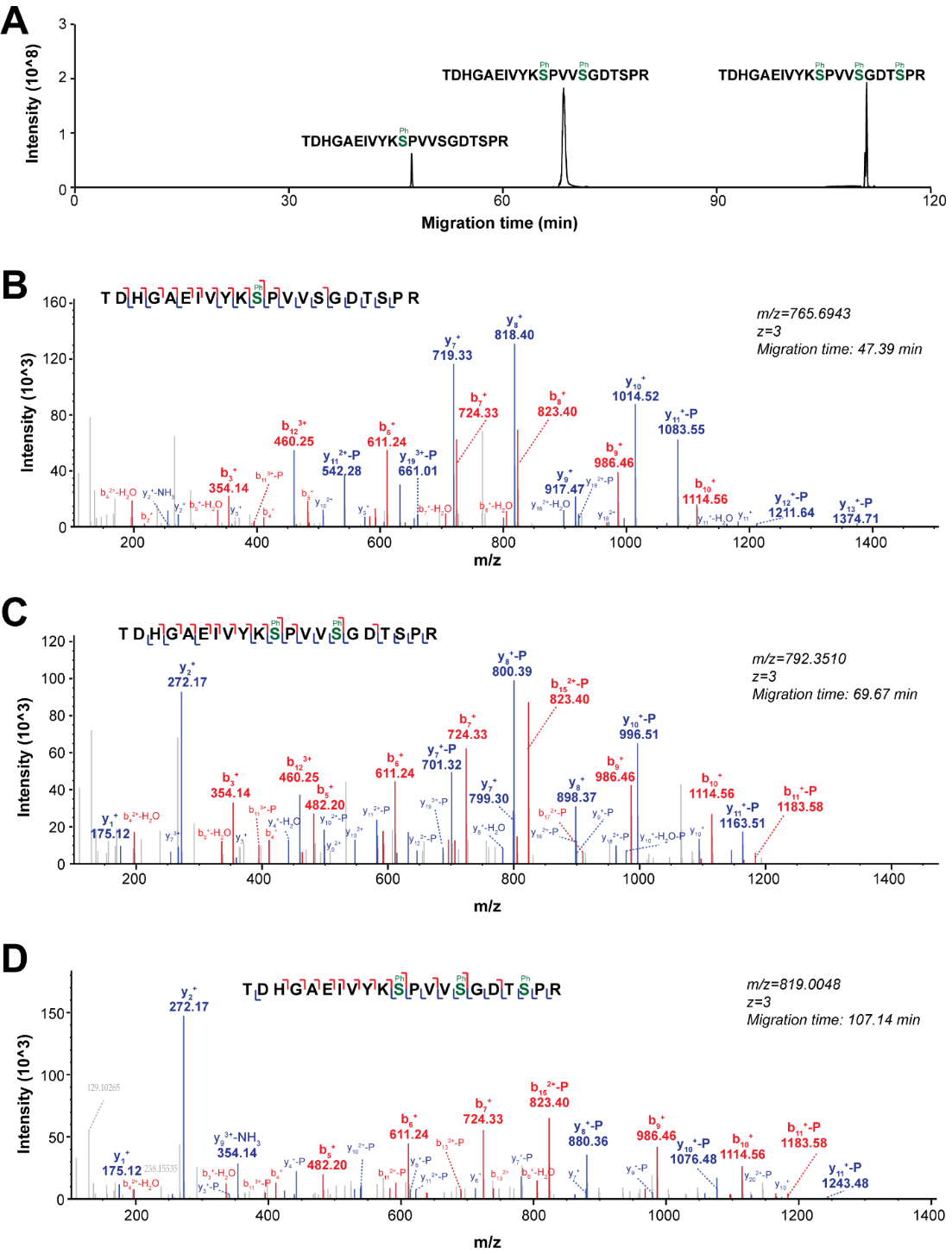

**Figure S2.** Example spectra of peptides carrying different numbers of phosphate groups. (A) Extracted ion electropherogram of peptide TDHGAEIVYKSPVVSGDTSPR with 1, 2, and 3 phosphorylation sites. Tandem mass spectra of peptide TDHGAEIVYKSPVVSGDTSPR carrying (B) 1, (C) 2, and (D) 3 phosphate groups.

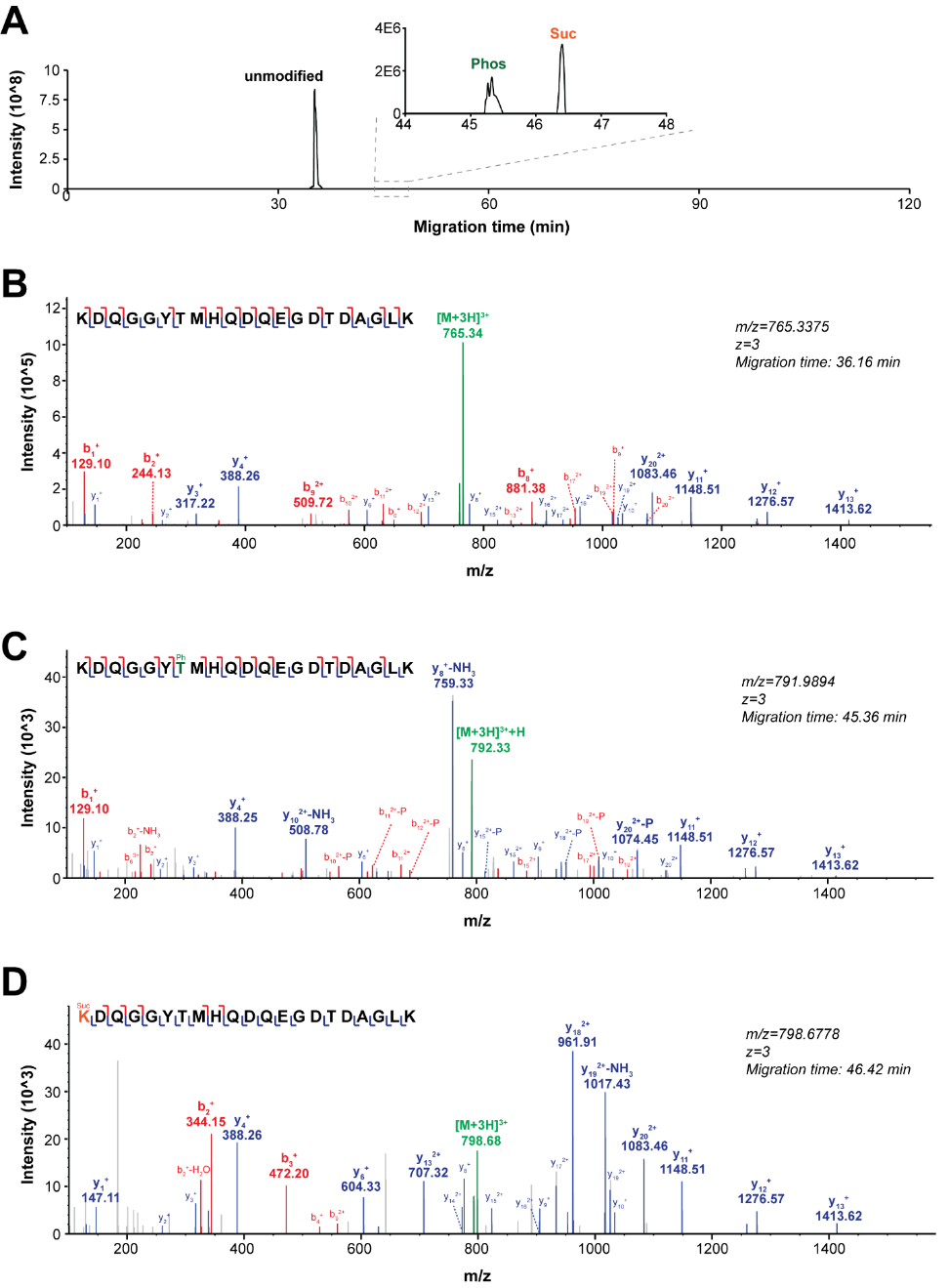

**Figure S3**. Example spectrum of peptide (KDQGGYTMHQDQEGDTDAGLK) without PTMs and with phosphorylation or succinylation. (A) Extracted ion electropherogram of the peptide without or with PTMs. Phos means phosphorylation and Suc indicates succinylation. (B)-(D) Annotated MS/MS spectra of the unmodified peptide (B), the phosphorylated peptide (C), and the succinylated peptide (D).

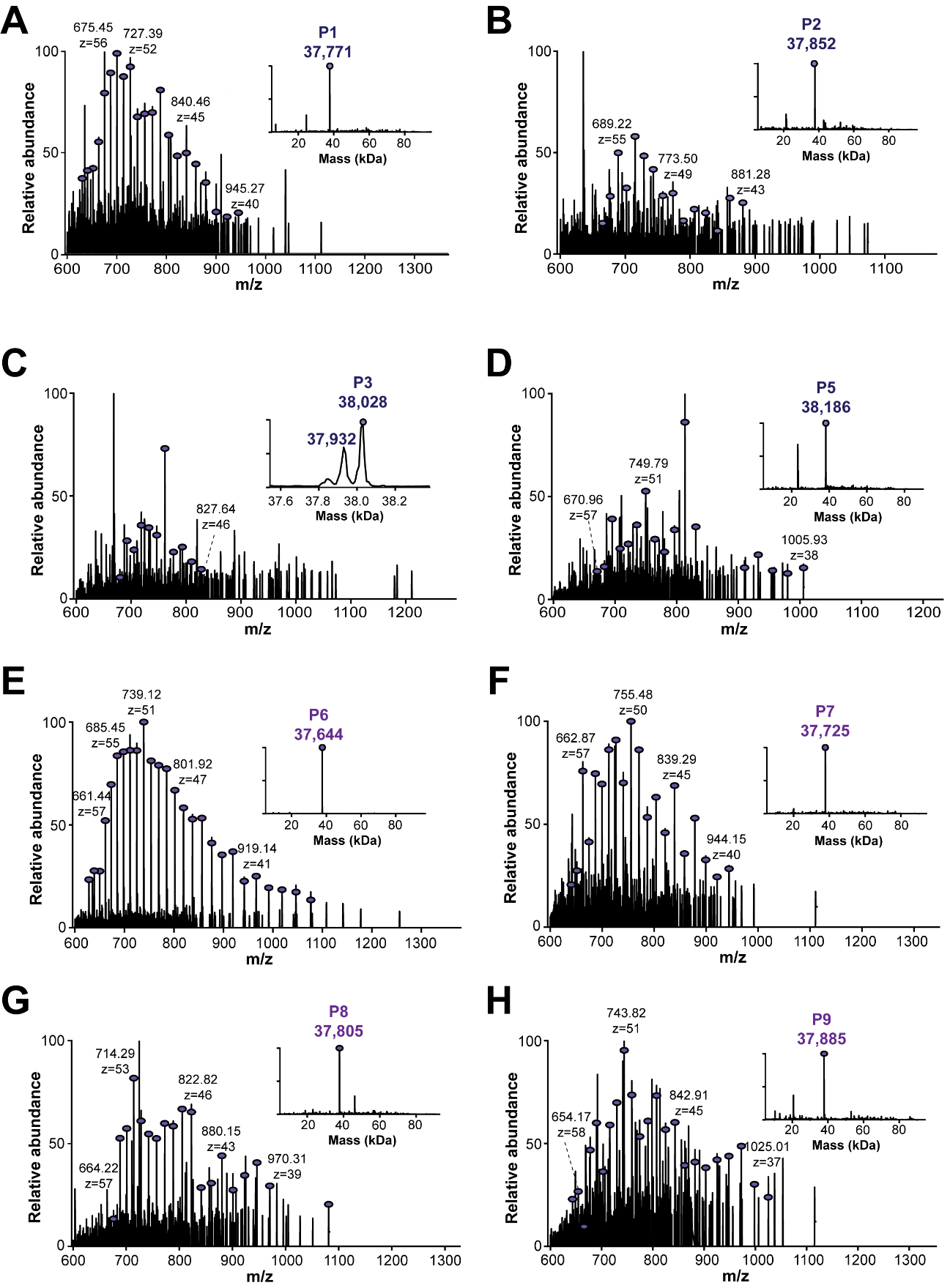

**Figure S4.** cIEF-MS analysis of human *p-tau-0N3R* under denaturing conditions. Mass spectra and deconvoluted masses of the proteoforms detected in P1 (A), P2 (B), P3 (C), P5 (D), P6 (E), P7 (F), P8 (G), and P9 (H).

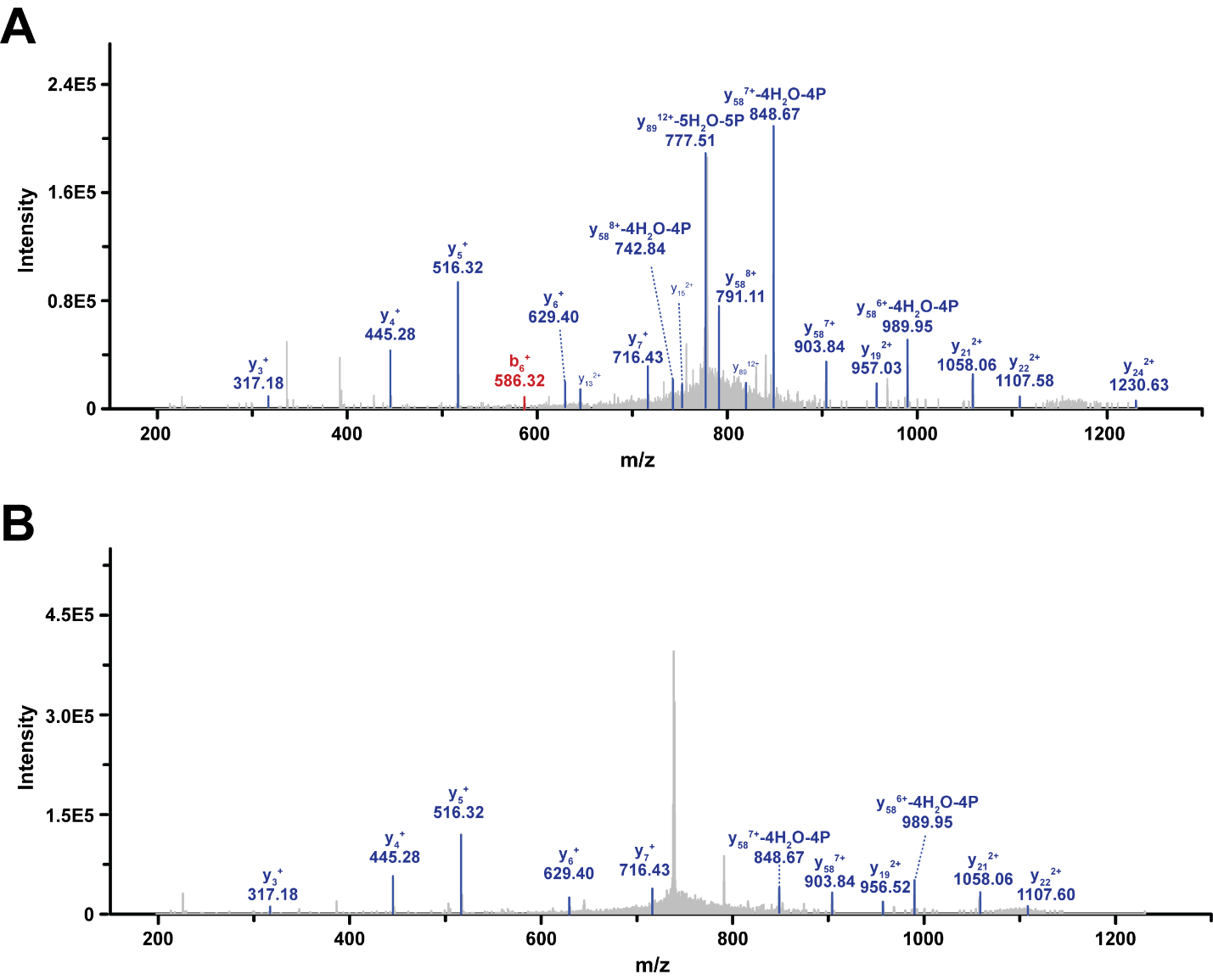

**Figure S5.** Annotated MS/MS spectra of two human *p-tau-0N3R* proteoforms in P4 (A) and P6 (B) carrying different numbers of phosphate groups under denaturing conditions.

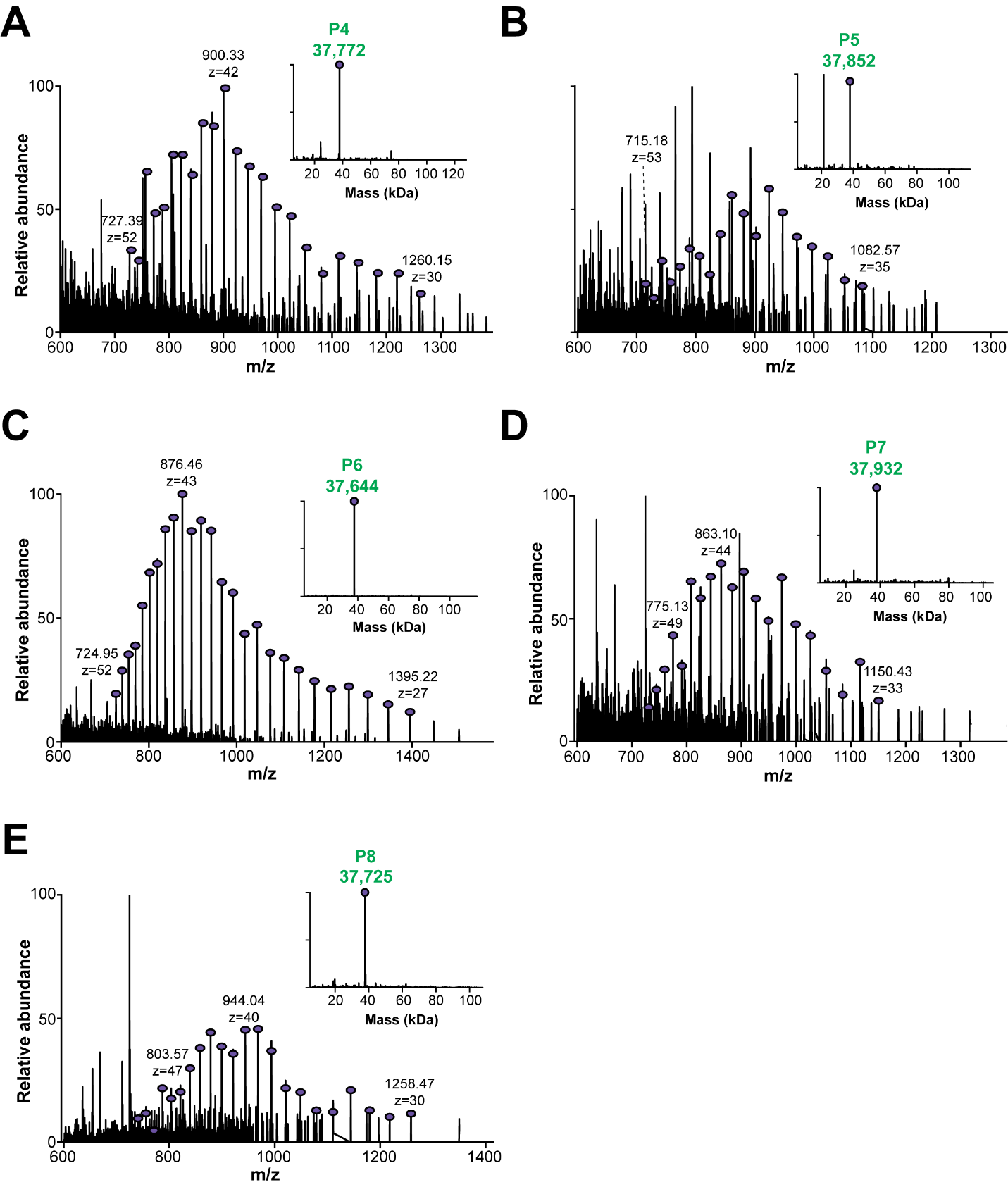

**Figure S6.** cIEF-MS analysis of human *p-tau-0N3R* under pseudo-native condition. (A-D) Averaged mass spectra and deconvoluted masses (inserted figures) of p-tau proteoforms detected in P4-P8.
